# RNA-EFM : Energy based Flow Matching for Protein-conditioned RNA Sequence-Structure Co-design

**DOI:** 10.1101/2025.02.09.637356

**Authors:** Abrar Rahman Abir, Liqing Zhang

**Author notes:** **Anonymous authors** Paper under double-blind review.

## Abstract

Ribonucleic acids (RNAs) are essential biomolecules involved in gene regulation and molecular recognition. Designing RNA molecules that can bind specific protein targets is crucial for therapeutic applications but remains challenging due to the structural flexibility of RNA and the laborious nature of experimental techniques. We propose **RNA-EFM**, a novel **E**nergy-based **F**low **M**atching framework for protein-conditioned RNA sequence and structure co-design. RNA-EFM integrates biophysical constraints, including the Lennard-Jones potential and sequence-derived free energy, to generate low-energy and biologically plausible RNA conformations. By incorporating an idempotent refinement strategy for iterative structural correction, RNA-EFM consistently outperforms existing baselines, achieving lower RMSD, higher lDDT, and superior sequence recovery across multiple evaluation splits.

## 1 Introduction

Ribonucleic acids (RNAs) are critical biomolecules that exhibit diverse functional roles in biological systems, including gene expression regulation, catalysis, and molecular binding. Their structural and functional versatility has paved the way for applications such as synthetic riboswitches for dynamic gene modulation and aptamers that target specific proteins (Dykstra et al., 2022; Thavarajah et al., 2021; Nori & Jin, 2024). However, experimental approaches such as SELEX, a high-throughput RNA selection technique, are often time-consuming and labor intensive, limiting their scalability for novel RNA therapeutics (Gold, 2015). This necessitates the development of computational frameworks that can efficiently design RNA sequences and structures to unlock their therapeutic potential (Sanchez de Groot et al., 2019).

Recent advancements in generative modeling have significantly impacted RNA design. Early methods for designing protein-binding RNAs focused on generating numerous candidate sequences followed by structural filtering or molecular dynamics simulations (Kim et al., 2010; Zhou et al., 2015; Buglak et al., 2020), which were computationally expensive and constrained by predefined structural motifs (Nori & Jin, 2024). Classical RNA structure design techniques such as algorithmic folding methods (Yesselman et al., 2019) have also been explored but lack scalability for complex sequence-structure co-design tasks. Modern generative approaches using diffusion models such as MMDiff (Morehead et al., 2023) have demonstrated improved performance for nucleic acid sequence-structure generation but often struggle with conditional generation and longer RNA sequences. Flow matching strategies, originally applied in protein structure design (Lipman et al., 2022; Bose et al., 2023), have shown promise for structure co-design tasks by interpolating between data distributions in a continuous manner. RNAFlow (Nori & Jin, 2024) builds on this by integrating an RNA inverse folding model with the pre-trained RF2NA network (Baek et al., 2024) for simultaneous sequence and structure generation. It models RNA conformational dynamics using inference trajectories, improving performance over standard methods. However, challenges remain in balancing efficiency and the accurate modeling of RNA’s dynamic structural flexibility, a critical factor in biological functionality (Ganser et al., 2019).

In this paper, we propose **RNA-EFM**, a novel framework for protein-conditioned RNA sequence-structure co-design. Our major contributions include: (1). **Protein-Conditioned RNA Sequence and Structure Co-Design:** RNA-EFM addresses the task of generating RNA sequences and structures jointly while being conditioned on protein interactions, enabling the design of functional RNAs tailored to specific protein targets. (2). **Energy-Based Flow Matching Framework:** We propose an energy-based flow matching framework that combines flow matching with an iterative refinement process based on the idempotent constraint, ensuring the generation of accurate RNA structures that progressively align better with the target. (3). **Incorporation of Biophysical Signals:** To further enhance structural quality and stability, we integrate biophysical constraints by adding the Lennard-Jones potential and sequence-derived free energy, guiding the model toward lower-energy RNA conformations while maintaining biological relevance. (4). **State-of-the-Art Performance:** RNA-EFM consistently surpasses existing baselines in both structure and sequence generation tasks, achieving lower RMSD, higher lDDT, and improved sequence recovery rates across multiple datasets.

## 2 Methods

### 2.1 Problem Formulation

RNA-EFM aims to generate RNA sequences and structures conditioned on protein backbone structures. Let the protein backbone atom structure be represented as 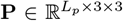, where *L*_*p*_ is the number of residues, and each residue contains backbone atoms *N, C*_*α*_, and *C*. The corresponding protein sequence is **p** = {*p*_*i*_ | *i* = 0, 1, …, *L*_*p*_ − 1}, where *p*_*i*_ is a categorical token. The RNA backbone structure is represented as 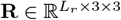 where *L*_*r*_ is the number of nucleotides, and each nucleotide contains backbone atoms P, 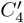, and *N*_1_/*N*_9_ (for pyrimidine and purine, respectively). The RNA sequence is **r** = {r_*i*_ |*i* = 0, 1, …, *L*_*r*_ −1}, where r_*i*_ is a nucleotide token. The task is to predict the RNA sequence **r** and backbone structure **R** conditioned on the protein structure **P** and sequence **p**.

### 2.2 Overview of the Approach

Our approach combines the principles of flow matching and energy-based refinement to generate biologically plausible RNA structures (Figure 1). The flow matching objective aligns the predicted structures with the target RNA backbone distribution, ensuring accurate geometric correspondence. Additionally, an iterative refinement process integrates biophysical energy constraints, leveraging a combination of sequence-derived free energy and atomic-level interactions. This approach allows the model to design RNA structures that are not only geometrically aligned with experimental targets but also energetically stable and biologically meaningful.

**Figure 1:**
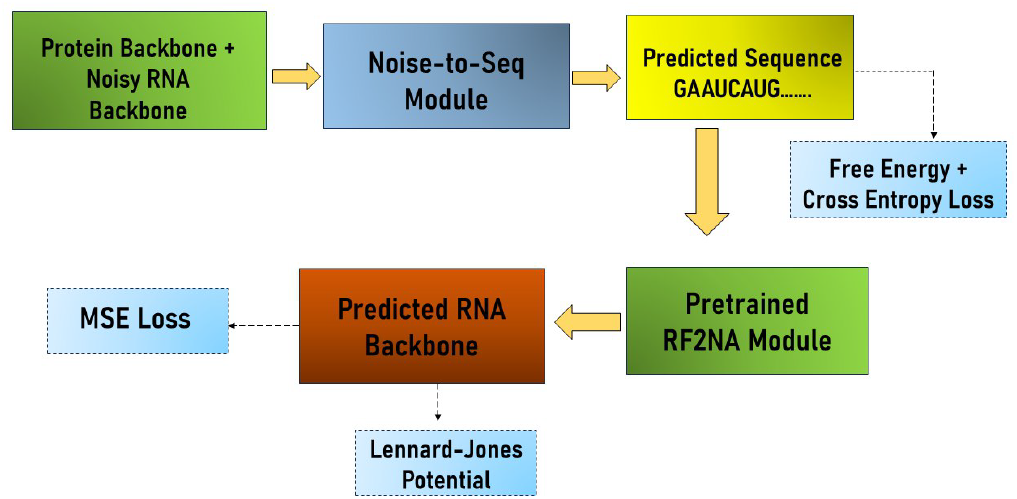
Overview of the RNA-EFM architecture. The model takes as input a protein backbone and a noisy RNA backbone. The Noise-to-Seq module predicts the RNA sequence, which is further processed using the pretrained RF2NA module to predict the RNA backbone structure. The predicted RNA backbone is refined iteratively by minimizing the Mean Squared Error (MSE) loss and incorporating the Lennard-Jones potential for structural stability. The predicted sequence is optimized using a combination of free energy and cross-entropy loss, ensuring biophysically plausible and structurally accurate RNA generation.

### 2.3 Flow Matching Objective

The RNA-EFM framework adopts flow matching to transform a prior distribution **p**_**0**_(**R**) into a target distribution **p**_**1**_(**R**), where **R** represents RNA backbone structures following (Nori & Jin, 2024). This transformation is achieved by parameterizing the flow as a sequence of time-dependent conditional probability distributions **p**_**t**_(**R**_**t**_|**R**_**1**_), where t ∈ [0, 1] denotes the interpolation step.

The intermediate RNA backbone structure **R**_**t**_ at any time step t is obtained through linear interpolation between a sample from the prior distribution **R**_**0**_ ∼ **p**_**0**_(**R**) and a sample from the target distribution **R**_**1**_∼ **p**_**1**_(**R**), given by **R**_**t**_ |**R**_**1**_ = (1− t)**R**_**0**_ + t**R**_**1**_, where t ∼ **U**(0, 1) is uniformly sampled. This interpolation constructs a continuous path from **p**_**0**_ to **p**_**1**_ that the model learns to approximate. To account for the geometric properties of RNA backbone structures, we utilize the Kabsch algorithm to align the prior sample **R**_**0**_ with the target structure **R**_**1**_. The Kabsch algorithm minimizes the root-mean-square deviation (RMSD) between two sets of points, ensuring invariance to rigid transformations such as rotation and translation. The aligned structure is given by **R**_**0**_^∗^ = **K**(**R**_**0**_, **R**_**1**_), where **K**(·,·) denotes the Kabsch alignment operation. The flow matching objective aims to minimize the discrepancy between the true and predicted vector fields that govern the transformation from **p**_**0**_ to **p**_**1**_. The true vector field **v**_**t**_(**R**_**t**_|**R**_**1**_) is defined as:

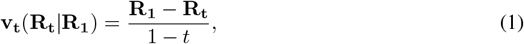

while the predicted vector field, parameterized by the neural network 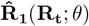 which predicts RNA backbone from a noisy intermediate backbone, is expressed as:

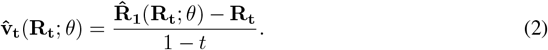

The flow matching loss **L**_**FM**_ is formulated as the expected squared difference between these vector fields:

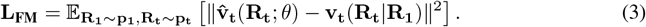

Substituting equation 1 and equation 2 into equation 3, we derive:

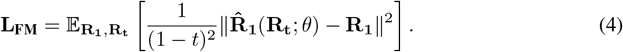

Finally, incorporating the Kabsch alignment to ensure invariance to rotational and translational transformations, the objective becomes:

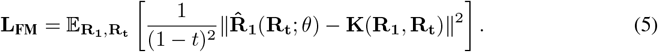

After marginalizing over t, the final loss reduces to the Mean Squared Error (MSE) between the aligned predicted and ground truth backbone structures:

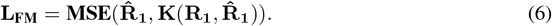

This formulation ensures that the RNA backbone predictions align accurately with the target structures while preserving geometric invariance.

## 3 Energy-Based Idempotent Flow Matching in RNA-EFM

To improve the biological plausibility of predicted RNA structures, RNA-EFM incorporates an energy-based refinement framework following (Zhou et al.) combining flow matching with biophysical constraints derived from RNA structure and sequence. This ensures that predicted structures align both geometrically and energetically with the target. The core idea is to iteratively refine the predicted RNA backbone structures by minimizing an energy function that penalizes deviations from the target structure while incorporating physical energy constraints for stability.

The refinement is governed by the conditional probability distribution:

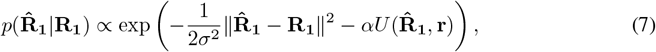

where 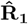 is the predicted RNA backbone structure, **R**_**1**_ is the target structure, and 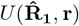 repre-sents the physical energy term combining the Lennard-Jones potential and the free energy computed from the predicted sequence using Vienna RNAfold (Lorenz et al., 2011). The Lennard-Jones potential is defined as:

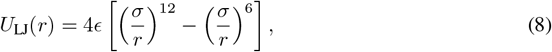

where r is the distance between two atoms, *σ* is the equilibrium distance where attractive and repulsive forces balance, and *ϵ* represents the depth of the potential well, determining interaction strength.

Taking the negative logarithm of the probability distribution in equation 7 results in the energy function:

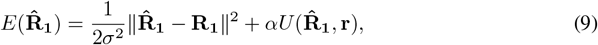

which is minimized during refinement. The gradient, used for iterative refinement, is given by:

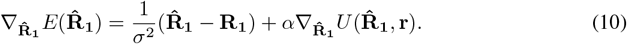

The refinement process is guided by the *idempotent flow matching property*, ensuring stabilization of predicted structures through repeated refinement until convergence. Mathematically, it is expressed as:

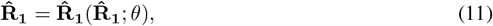

indicating the predictor converges to a fixed point on the manifold of stable RNA structures when applied iteratively. The training objective incorporates both flow matching and biophysical energy constraints, defined as:

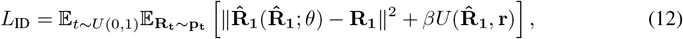

where the first term minimizes deviation from the target structure while the second term penalizes high-energy configurations. Minimizing this objective ensures RNA-EFM generates both geometrically accurate and energetically favorable RNA structures. See Appendix A for detailed training process.

## 4 Experiments

### 4.1 Dataset

We use the filtered PDBBind dataset for training and evaluation. Protein-RNA complexes are filtered to ensure at least one protein Cα atom and RNA C’4 atom are within 7 Å, following (Liu et al., 2017). RNA chains are restricted to lengths between 6 and 96, and protein chains are cropped to a contiguous length of 50. All experiments were conducted on two splits-sequence similarity split and RF2NA (RoseTTAFold2NA (Baek et al., 2024)) split. For the sequence similarity split, the dataset includes 1015 train, 105 validation, and 72 test complexes. The RF2NA split comprises 1059 train, 117 validation, and 16 test complexes.

### 4.2 Baselines

To evaluate RNA-EFM, we compare it against multiple baselines for RNA structure and sequence generation. For structure generation, we consider Conditional MMDiff (Morehead et al., 2023), an SE(3)-equivariant diffusion model, and RNAFlow (Nori & Jin, 2024), a flow matching-based framework, along with its variants. RNAFlow-Base, initializes structure generation using RF2NA by folding a mock RNA sequence composed entirely of adenines and iteratively refining the predicted conformation, while RNAFlow-Traj conditions on multiple RNA backbone conformations. RNAFlow-Base + Rescore and RNAFlow-Traj + Rescore further enhance selection through a rescoring model. For sequence generation, we include a Random baseline, which selects nucleotides uniformly, an LSTM-based model that autoregressively predicts RNA sequences (Im et al., 2019), Conditional MMDiff and RNAFlow along with its variants.

## 5 Results

### Metrics

We evaluate structure generation using RMSD (aligned with the Kabsch algorithm) and lDDT, both capturing structural accuracy. For sequence generation, we report recovery rate, the percentage of correctly predicted nucleotides.

As shown in Tables 1 and 2, RNA-EFM consistently outperforms all baselines on both structure and sequence generation tasks. For structure generation, RNA-EFM achieves a 5.75% reduction in RMSD and a 13.21% improvement in lDDT on the RF2NA split compared to the best baseline. On the sequence similarity split, RNA-EFM demonstrates a 10.96% reduction in RMSD and a 5.26% improvement in lDDT. For sequence recovery, RNA-EFM shows an 8.11% improvement on the RF2NA split and a 9.37% improvement on the sequence similarity split compared to the best-performing RNAFlow baseline. These results validate the effectiveness of RNA-EFM’s energy-based refinement and idempotent flow matching framework in generating accurate and biologically plausible RNA structures and sequences.

**Table 1:** RNA structure generation results. We report Mean ± SEM for RMSD and IDDT metrics.

| Method | RF2NA Pre-Training Split |  | Sequence Similarity Split |  |
| --- | --- | --- | --- | --- |
|  | RMSD | IDDT | RMSD | IDDT |
| Conditional MMDiff | 14.82 ± 1.01 | 0.34 ± 0.02 | 17.42 ± 0.86 | 0.38 ± 0.01 |
| RNAFlow-Base | 12.85 ± 0.63 | 0.51 ± 0.01 | 14.77 ± 0.34 | 0.57 ± 0.01 |
| RNAFlow-Traj | 13.12 ± 0.64 | 0.52 ± 0.01 | 15.11 ± 0.33 | 0.57 ± 0.00 |
| RNAFlow-Base + Rescore | 10.61 ± 1.73 | 0.53 ± 0.03 | 14.60 ± 1.05 | 0.56 ± 0.02 |
| RNAFlow-Traj + Rescore | 15.30 ± 1.89 | 0.52 ± 0.03 | 15.31 ± 0.93 | 0.56 ± 0.02 |
| **RNA-EFM** | **10.00 ± 0.50** | **0.60 ± 0.01** | **13.00 ± 0.40** | **0.60 ± 0.01** |

**Table 2:** RNA sequence generation results. We report Mean ± SEM for native sequence recovery.

| Method | RF2NA Pre-Training Split | Sequence Similarity Split |
| --- | --- | --- |
| Random | 0.25 ± 0.00 | 0.25 ± 0.00 |
| LSTM | 0.27 ± 0.01 | 0.24 ± 0.01 |
| Conditional MMDiff | 0.24 ± 0.02 | 0.22 ± 0.02 |
| RNAFlow-Base | 0.30 ± 0.02 | 0.30 ± 0.01 |
| RNAFlow-Traj | 0.31 ± 0.01 | 0.28 ± 0.01 |
| RNAFlow-Base + Rescore | 0.33 ± 0.02 | 0.32 ± 0.03 |
| RNAFlow-Traj + Rescore | 0.37 ± 0.05 | 0.29 ± 0.02 |
| **RNA-EFM** | **0.40 ± 0.03** | **0.35 ± 0.02** |

## 6 Conclusion

In this work, we presented RNA-EFM, a framework for RNA sequence and structure generation conditioned on protein interactions. By integrating biophysical energy-based refinement with idempotent flow matching, RNA-EFM ensures the generation of geometrically accurate, energetically stable, and biologically relevant RNA structures. Our results show that RNA-EFM significantly outperforms baselines in both structure and sequence generation tasks, demonstrating its potential to advance the design of functional RNAs tailored to protein targets.

## Appendix

### A Training and Inference in RNA-EFM

The idempotent objective in RNA-EFM facilitates iterative refinement of the predicted RNA backbone structures, ensuring convergence to stable and biologically plausible configurations. While theoretically, the structure predictor 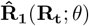 could refine the predicted structure an infinite number of times, excessive iterations would lead to increased computational overhead during inference. To address this, we restrict the refinement process to a single step per iteration during training and inference, balancing computational efficiency with refinement quality.

In our implementation, following Zhou et al., the refinement framework employs a predictor-refiner strategy. At each step, the structure predictor generates an initial prediction 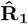, which is then refined to align with the target structure **R**_**1**_. This approach aligns well with the idempotent property, ensuring that further refinements stabilize the predicted structure. During training, comprising 50% of the total iterations, the structure predictor is optimized using the flow matching objective. This phase ensures that the predictor aligns the predicted backbone structures with the target structures. In the subsequent 50% of training, the refinement process explicitly incorporates energy based biophysical constraints, guided by the idempotent loss function defined in equation 12. However, following Nori & Jin (2024), we employ **Noise-to-seq** + RF2NA (C) as our predictor where Noise-to-seq predicts the sequence and pre-trained RF2NA (RoseTTAFold2NA) predicts the backbone. We also add cross entropy loss between predicted and true nucleotides. We follow the similar inference algorithm like Nori & Jin (2024). At inference time, the structure predictor outputs refined structures that conform to both the target geometry and energetically favorable configurations, as defined by the energy-based refinement framework. Algorithm B describes the training process.

### B Algorithm

#### Algorithm 1

RNA-EFM Training Algorithm

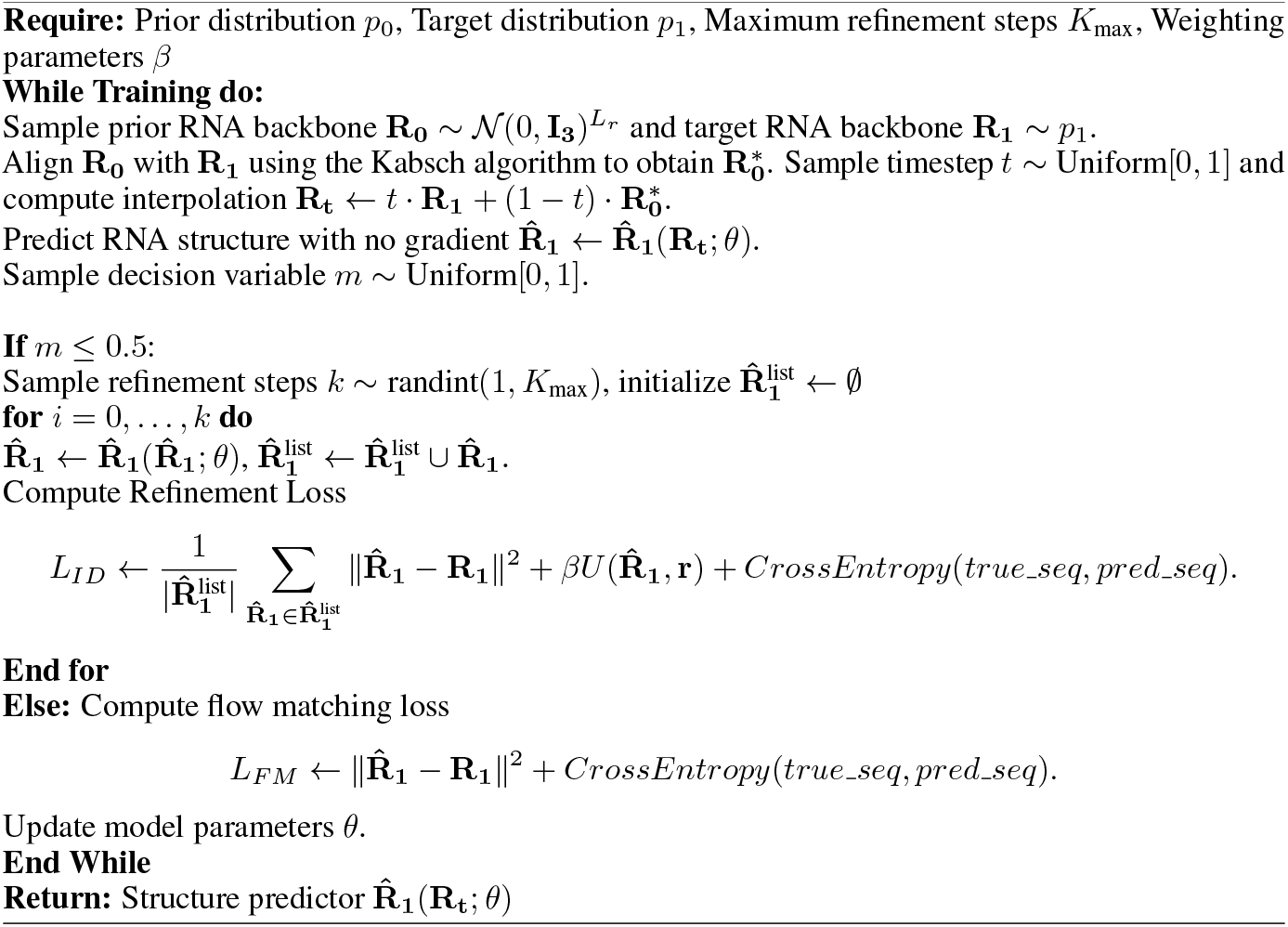

### C Noise-To-Seq Module

We describe the Noise-to-Seq from Nori & Jin (2024) for completeness. Noise-to-Seq is a graphbased RNA inverse folding model, to predict RNA sequences autoregressively from noised structures. The model uses an encoder-decoder architecture where the encoder processes a protein-RNA complex and the decoder predicts a probability distribution over nucleotides for sequence generation.

#### Graph Representation

Each 3D backbone point cloud 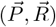 is represented as a graph G = (𝒱, ℰ). Nodes 𝒱 include amino acids at the *C*_*α*_ coordinates of the protein 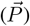 and nucleotides at the 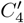 coordinates of the RNA 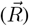. For each node 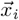, edges ℰ are drawn to its 10 nearest neighbors within the same graph and to the 5 nearest neighbors across graphs. Edge features are defined by the Euclidean distance:

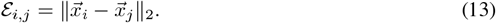

#### Node and Edge Features

Node features include unit vectors to neighboring nodes, residue type (nucleotide or amino acid), and local backbone orientation. Edge features include magnitude and directional information, as well as indicators for cross-graph edges.

#### Model Architecture

The encoder applies Geometric Vector Perceptrons (GVPs) to both node and edge features *υ*_*i*_ and ℰ_*i,j*_:

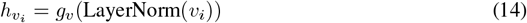

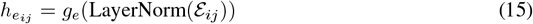

where g_*υ*_ and g_*e*_ denote the GVP-based embeddings. Node embeddings are updated using a sequence of three message-passing GVP layers followed by a residual connection:

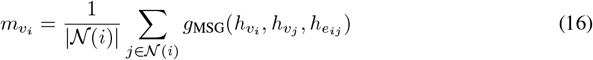

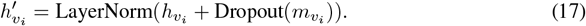

where g_MSG_ denotes a sequence of GVP layers and 𝒩 (*i*) is the set of neighbors for node *i*. During sequence generation, an additional timestep embedding is included. The decoder autoregressively predicts nucleotides using a softmax layer, and the model is supervised using a cross-entropy loss function comparing the predicted and true nucleotide classes:

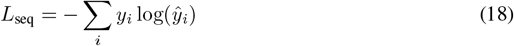

where *y*_*i*_ is the true nucleotide label and 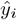 is the predicted probability for position *i*.

## References

Minkyung Baek, Ryan McHugh, Ivan Anishchenko, Hanlun Jiang, David Baker, and Frank DiMaio. Accurate prediction of protein–nucleic acid complexes using rosettafoldna. Nature methods, 21(1): 117–121, 2024.

Avishek Joey Bose, Tara Akhound-Sadegh, Guillaume Huguet, Kilian Fatras, Jarrid Rector-Brooks, Cheng-Hao Liu, Andrei Cristian Nica, Maksym Korablyov, Michael Bronstein, and Alexander Tong. Se (3)-stochastic flow matching for protein backbone generation. arXiv preprint arXiv:2310.02391, 2023.

Andrey A Buglak, Alexey V Samokhvalov, Anatoly V Zherdev, and Boris B Dzantiev. Methods and applications of in silico aptamer design and modeling. International journal of molecular sciences 21(22):8420, 2020.

Peter B Dykstra, Matias Kaplan, and Christina D Smolke. Engineering synthetic rna devices for cell control. Nature Reviews Genetics 23(4):215–228, 2022.

Laura R Ganser, Megan L Kelly, Daniel Herschlag, and Hashim M Al-Hashimi. The roles of structural dynamics in the cellular functions of rnas. Nature reviews Molecular cell biology 20(8):474–489, 2019.

Larry Gold. Selex: How it happened and where it will go. Journal of molecular evolution, 81(5): 140–143, 2015.

Jinho Im, Byungkyu Park, and Kyungsook Han. A generative model for constructing nucleic acid sequences binding to a protein. BMC genomics, 20(Suppl 13):967, 2019.

Namhee Kim, Joseph A Izzo, Shereef Elmetwaly, Hin Hark Gan, and Tamar Schlick. Computational generation and screening of rna motifs in large nucleotide sequence pools. Nucleic acids research 38(13):e139–e139, 2010.

Yaron Lipman, Ricky TQ Chen, Heli Ben-Hamu, Maximilian Nickel, and Matt Le. Flow matching for generative modeling. arXiv preprint arXiv:2210.02747, 2022.

Zhihai Liu, Minyi Su, Li Han, Jie Liu, Qifan Yang, Yan Li, and Renxiao Wang. Forging the basis for developing protein–ligand interaction scoring functions. Accounts of chemical research, 50(2): 302–309, 2017.

Ronny Lorenz, Stephan H Bernhart, Christian Höner zu Siederdissen, Hakim Tafer, Christoph Flamm, Peter F Stadler, and Ivo L Hofacker. Viennarna package 2.0. Algorithms for molecular biology, 6: 1–14, 2011.

Alex Morehead, Jeffrey Ruffolo, Aadyot Bhatnagar, and Ali Madani. Towards joint sequence-structure generation of nucleic acid and protein complexes with se (3)-discrete diffusion. arXiv preprint arXiv:2401.06151, 2023.

Divya Nori and Wengong Jin. Rnaflow: Rna structure & sequence design via inverse folding-based flow matching. arXiv preprint arXiv:2405.18768, 2024.

Natalia Sanchez de Groot, Alexandros Armaos, Ricardo Graña-Montes, Marion Alriquet, Giulia Calloni, R Martin Vabulas, and Gian Gaetano Tartaglia. Rna structure drives interaction with proteins. Nature communications 10(1):3246, 2019.

Walter Thavarajah, Laura M Hertz, David Z Bushhouse, Chloé M Archuleta, and Julius B Lucks. Rna engineering for public health: innovations in rna-based diagnostics and therapeutics. Annual review of chemical and biomolecular engineering 12(1):263–286, 2021.

Joseph D Yesselman, Daniel Eiler, Erik D Carlson, Michael R Gotrik, Anne E d’Aquino, Alexandra N Ooms, Wipapat Kladwang, Paul D Carlson, Xuesong Shi, David A Costantino, et al. Computational design of three-dimensional rna structure and function. Nature nanotechnology 14(9):866–873, 2019.

Qingtong Zhou, Xiaole Xia, Zhaofeng Luo, Haojun Liang, and Eugene Shakhnovich. Searching the sequence space for potent aptamers using selex in silico. Journal of chemical theory and computation 11(12):5939–5946, 2015.

Wenyin Zhou, Christopher Iliffe Sprague, and Hossein Azizpour. Energy-based flow matching for molecular docking.

